# Synthetic memory circuits for programmable cell reconfiguration in plants

**DOI:** 10.1101/2022.02.11.480167

**Authors:** James P B Lloyd, Florence Ly, Patrick Gong, Jahnvi Pflüger, Tessa Swain, Christian Pflüger, Muhammad Adil Khan, Brendan Kidd, Ryan Lister

## Abstract

Plant biotechnology predominantly relies on a restricted set of genetic parts with limited capability to customize spatiotemporal and conditional expression patterns. Synthetic gene circuits have the ability to integrate multiple customizable input signals through a processing unit constructed from biological parts, to produce a predictable and programmable output. Here, we present a suite of functional recombinase-based gene circuits for use in plants. We first established a range of key gene circuit components compatible with plant cell functionality. We then used these to develop a range of operational logic gates using the identify function (activation) and negation function (repression) in Arabidopsis protoplasts and *in vivo*, demonstrating their utility for programmable manipulation of transcriptional activity in a complex multicellular organism. Through utilization of genetic recombination these circuits create stable long-term changes in expression and recording of past stimuli. This highly-compact programmable gene circuit platform provides new capabilities for engineering sophisticated transcriptional programs and previously unrealised traits into plants.

## Introduction

A great deal of success has been made in plant biotechnology with relatively simple genetic tools, such as strong constitutive promoters. However, continuous overexpression of a gene across a whole plant can be detrimental to its growth patterns^1–5^. Natural gene regulatory systems can integrate multiple signals to activate or repress transcriptional output, but currently few tools allow sophisticated spatiotemporal control of transgenes in plants. Synthetic gene circuits are a promising approach to overcome this limitation^6,7^. Ideally, gene circuits would take multiple input signals and integrate them into a genetic processing unit to control an output gene’s expression in a user-defined manner. These circuits aim to function in a manner analogous to natural gene regulatory networks, which can be extremely complex, with multiple inputs, outputs and cross-talk between factors^8,9^. Some natural gene regulatory units closely resemble simple Boolean logic gates. For example, the *lac* operon in *E. coli* approximates an A NIMPLY B Boolean logic gate^10,11^, with lactose activating expression but glucose repressing expression, overriding the presence of lactose^10,11^. Engineering and application of synthetic logic gates in plants could provide us with the ability to tailor cellular activity and growth, programming them with sophisticated traits that cannot be achieved by conventional transcriptional control technologies.

In bacteria, yeasts, and mammalian cells, transcription factors and recombinases have been used to construct gene circuits in cell culture^12–23^. However, in plants, only a limited number of circuits have been developed to date. These include the split-TALE to act as an AND gate^24^, the recombinase-based toggle switch^25^ and an elegant system using a combination of bacterial transcription factors to generate various types of logic^7^. While important advances, these circuit designs are however either limited in their range of logic and application, or have output activity that is coupled to the persistent presence of the input signal(s). Alternative memory gene circuit technologies that drive output activity that persists beyond input signal presence are therefore needed for applications that are dependent on long-lived changes in gene expression.

Cell fate decisions in plant development require a cellular memory of specific experiences^26^. This is challenging to replicate in synthetic gene circuits, as the effect is often transient due to the targeted expression change being temporally linked to the inducing signal. However, with recombination-based gene circuits, an altered expression pattern and resulting cell state can be locked in. Rather than doing this with epigenetic changes, as done in nature^26^, recombination based circuits achieve this by permanent changes to the genome in a specific subset of cells^19^. Therefore this system is more akin to the adaptive immune system of vertebrates that creates unique T-cell receptors and antibodies via recombination^27^, than a traditional cell fate mechanism that can be reversed^28^. Recombinase-based gene circuits can therefore take an analogue input signal and convert it into a digital signal^20^.

In mathematical terminology, the identify function describes what happens when an input being on causes the output to switch on. In terms of gene expression, this is gene activation from a positive signal. In contrast, the mathematical negation (NOT) function would be equivalent to gene repression from a negative signal. To build all of the basic Boolean logic gates as gene circuits, both the identify and negation functions need to be implemented using biological parts. The ‘Boolean logic and arithmetic through DNA excision’ (BLADE) system, developed in mammalian cell culture systems^19^, uses recombinases to either activate a gene by removing a terminator sequence between the promoter and coding sequence of the output gene, or repress gene expression by removing a part of the output gene^19^. Such a recombinase-based design means that the switch from one transcriptional state to another is long-term and stable, and thus the continuous addition of the activating signal is not required, unlike gene circuit designs based on transcription factors. Given the limited range of plant circuits for the control of plant cellular activity and development, especially with respect to circuits able to generate long-term stable changes in gene expression, here we aimed to develop, optimize, and implement effective recombinase-based synthetic gene circuits to enable new capabilities for sophisticated plant engineering that requires memory-based functions.

## Results

To create recombinase-based gene circuits with the identify function, we need to repress output gene expression until activated. In mammalian and bacterial recombinase-based gene circuits^19,21^, repression has been achieved by placement of transcriptional terminators between the promoter and coding DNA sequence (CDS) of a gene. When flanked by recombination sites the terminator can be removed upon recombinase expression, however, the transcriptional terminators used in these studies are unlikely to be effective across kingdoms. Fortunately, multiple plant terminators have previously been characterised^29–31^ and used to block recombinase activity during lineage tracing in Arabidopsis roots^32^. To find a highly repressive terminator for our circuit design, we used a high-throughput dual luciferase assay to measure the output from constructs transfected into Arabidopsis leaf mesophyll protoplasts^33^. Each construct contained both a *Renilla luciferase* (*Rluc*) gene as the circuit output reporter, and a *Firefly luciferase* (*Fluc*) gene as the normaliser to account for variation in transfection efficiency (Fig. 1a). The ratio of Rluc to Fluc activity reflects the gene circuit output activity. We compared the ability of multiple candidate repressive sequences to inhibit *Rluc* reporter expression (Fig. 1b): (1) the short synthetic *transcription blocker* (*TB*)^34^; (2) *Octopine Synthase* (*OCS)* terminator from *Agrobacterium tumefaciens*^29,32^; (3) *OCS* followed by *TB*, as stacking terminators can often enhance activity^30^; (4) a variant of the *OCS* terminator^35^; (5) the *35S* cauliflower mosaic virus (CaMV) terminator^29^; and (6) an upstream open reading frame (uORF) from *AT1G36730*, being a conserved post-transcriptional repressor^36–38^. All of these plasmids were compared to a constitutively active construct that lacks the addition of any repressor or any recombination sites (Fig. 1b). The *FRT* recombination sites recognized by the Flp recombinases were incorporated to flank the candidate repressive sequences. All versions of the *OCS* terminator were able to repress target gene expression 24 hours after construct transfection into Arabidopsis protoplasts (Fig. 1c). Only mild repression was observed for the *TB* and uORF, and no repression was observed with the *CaMV* terminator (Fig. 1c), potentially due to previously observed functional attenuation when the *CaMV* terminator is placed close to the promoter^39^. Therefore, all subsequent experiments to develop functional gene circuits used the *OCS* terminator to repress reporter expression. Next, we needed to identify a promoter that would drive constitutive recombinase expression at a level sufficient to fully activate the *Rluc* reporter gene by removing the *OCS* terminator. We tested the *NOS, 35S, Act2*, and *TCTP1* promoters^29,40^, which are commonly used constitutive plant promoters. Twenty-four hours after construct transfection into Arabidopsis protoplasts, all tested promoters increased reporter expression compared to the *OCS* terminator (Fig. 1d), but only the *TCTP1* promoter resulted in reporter expression levels comparable to the constitutive Rluc control (Fig. 1d). When constructing these plasmids, either an empty end-linker or one with the *lacZ* gene were cloned, with the latter allowing for selection of correct assembly^29,41^. To rule out that this can alter the expression of the construct, we tested *TCTP1::Flp* with and without the *lacZ* end-linker and observed no difference (Supp. Fig. 1; p = 1.0, Tukey HSD *post-hoc* test). To test whether the recombinase circuits function in whole plants, we generated stable transgenic Arabidopsis plants containing nuclear GFP driven by the *Act2* promoter and blocked by the *OCS* terminator alone (Fig. 1e). Expression of the nuclear GFP was only observed after induction of expression of the Flp recombinase, driven by the heat-activated by the *HSP18*.*2* promoter, by heat-shock at 37 °C for 5 hours (Fig. 1e), demonstrating that our gene circuit design can work *in vivo* with a condition-specific promoter driving the recombinase. Thus, here we demonstrate establishment of all the basic components required to develop recombination-based gene circuits: a strong terminator (*OCS*) to repress expression of the output gene, promoters able to drive expression of the recombinase to activate the circuit (*TCTP1* and *HSP18*.*2*), and effective recombination activity using the Flp recombinase and its FRT recombination sites.

**Fig. 1:**
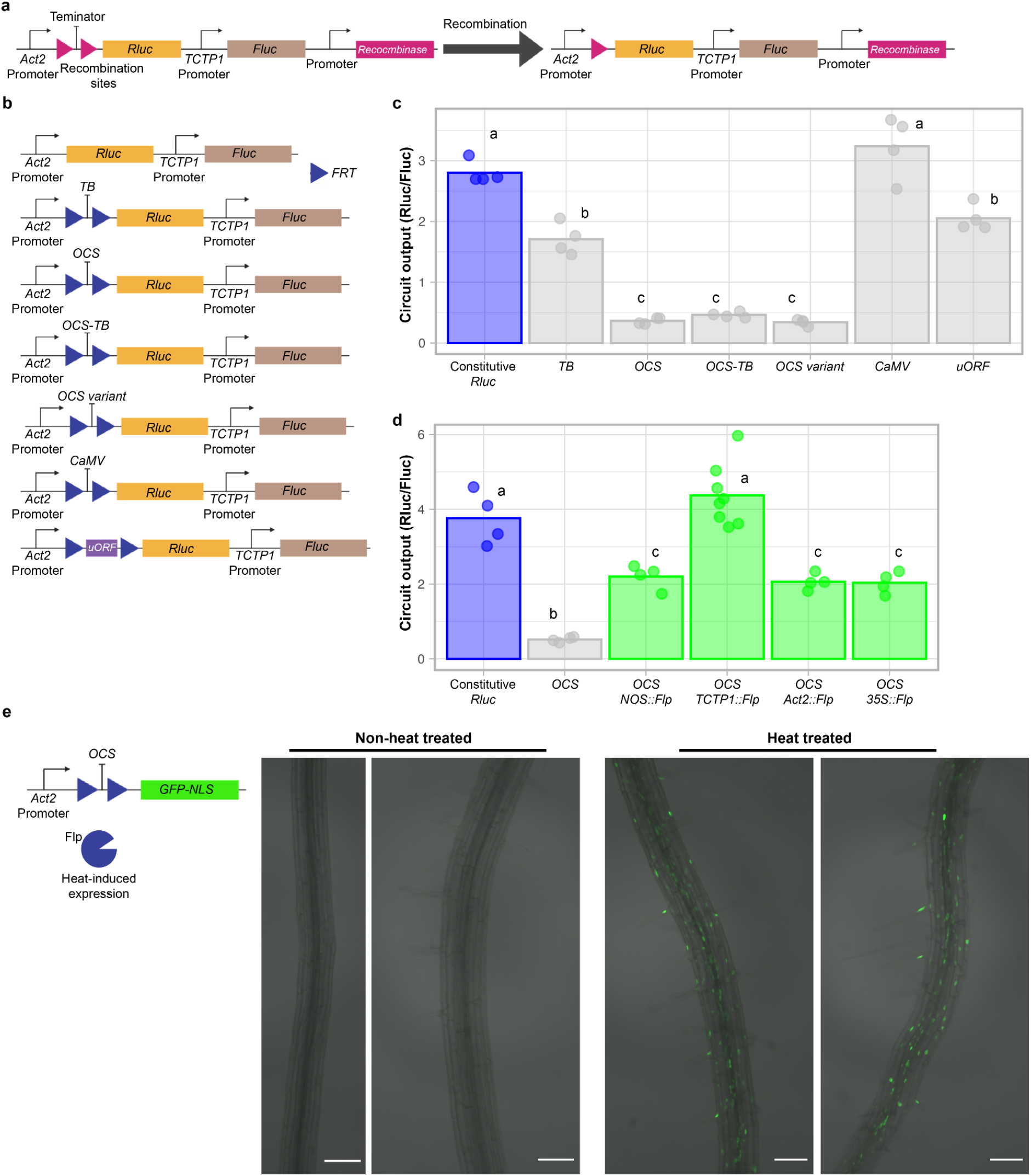
Recombination-based gene circuits built with plant parts can control plant gene expression. **a**, Schematic of recombination-based identify gene circuit construct design, before and after recombination has occurred. **b**, Constructs to identify a repressor of gene expression. The *Act2* promoter followed by the TMV 5’ UTR was used to drive the output gene (*Rluc* or *GFP*) and the *TCTP1* promoter was used to drive *Fluc*, which was used to normalise for transfection efficiency. The test constructs each had either a terminator of transcription or a uORF flanked by FRT sites, upstream of the *Rluc* coding sequence within the TMV 5’ UTR. **c**, Test of a range of terminators or uORFs, flanked by *FRT* recombination sites, upstream of *Rluc* reporter gene. Circuit output activity is the Rluc / Fluc luminescence ratio; also for panel d) 24 hours after protoplast transfection (n=4); Crossbar, mean. The blue bar is the control samples, gray bars represent samples that are expected to be repressed, and bars with different letters have a significant difference calculated by a one-way ANOVA and Tukey’s HSD test (p < 0.05) (also for panel d). **d**, Test of different promoters driving *Flp* recombinase expression 24 hours after transfection into Arabidopsis protoplasts (n=4). Crossbar, mean. Bar colours as per panel c, with green bars representing samples expected to be activated. **e**, Induction of nuclear localised GFP in Arabidopsis roots after heat-shock in transgenic Arabidopsis plants harboring the *HSP18*.*2* promoter driving heat-dependent *Flp* expression to remove the *OCS* terminator from the *Act2* promoter that blocks *GFP* transcription. The image is merged from transmitted light and GFP channels and is the max intensity projection from 10 slices. Scale bar = 100 μm. Some figure components created with BioRender.

To develop 2-input Boolean logic gates in plants, a second recombinase is required to act as the second input. Initially, we assembled and tested an OR gate using Cre as the second recombinase, which has previously been used in Arabidopsis^32,42–44^. Surprisingly, we found that Cre was unable to activate an OR gate when expressed alone in Arabidopsis protoplasts (Fig. 2a). Furthermore, when Cre and Flp were both expressed, the robust activation seen when Flp was expressed alone was inhibited (Fig. 2). To confirm that our construct could be recognized and recombined by Cre, we used purified Cre to cut the recombination site of a purified OR gate plasmid *in vitro* and confirmed that recombination took place (Supp. Fig. 2a). This suggests that the Cre protein was inhibitory to circuit function in our system. Therefore, we tested four more recombinases (B3, SCre, VCre, Vika) to find one that could act as a second input (Fig. 2b and c and Supp. Fig. 2d). Of the four tested, only B3 led to strong activation, comparable to the activation produced by Flp (Fig. 2c and Supp. Fig. 2d). To confirm that there is no cross-reactivity between Flp and B3, we compared the expression of our reporter when B3 and Flp recombinases were paired with their non-cognate recombination sites (Fig. 2d). No cross-reactivity was observed (Fig. 2e), but there was a high rate of reporter activation when recombinases were paired with their cognate recombination site. Taken together, this identifies B3, but not Cre, as a suitable secondary recombinase to act alongside Flp in dual input circuits.

**Fig. 2:**
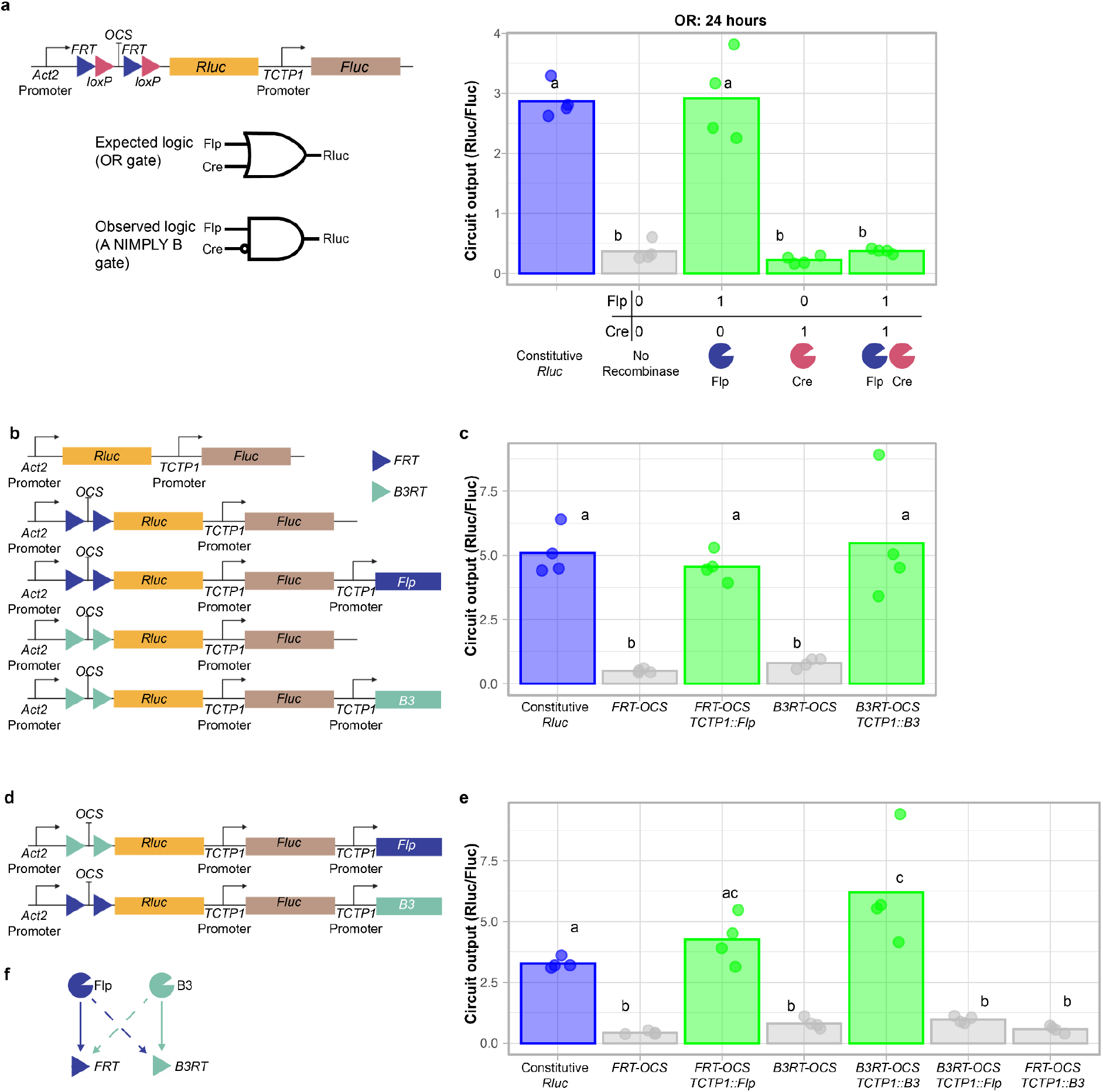
Identification of a additional functional recombinases for the construction of complex gene circuits. **a**, Schematic of the Cre/Flp-based OR gate construct design and the Cre/Flp-based OR gate that produces high circuit output activity only when Flp is present alone, 24 hours post-transfection (as in panels c and e). Circuit output activity is the Rluc / Fluc luminescence ratio; also for panel d) 24 hours after protoplast transfection (n=4); Crossbar, mean. Blue bar represent the control sample, green bars represent samples expected to be activated (turned on), gray bars represent samples expected to be repressed (turned off) and bars with different letters have a significant difference calculated by a one-way ANOVA and Tukey’s HSD test (p < 0.05) (also for panel c and e). **b**, Constructs used to test whether the B3 recombinase can adequately remove the OCS terminator and activate Rluc expression in plant cells, compared to Flp. **c**, Test of B3 for identify (activation) function in plant cells, compared to the previously tested Flp-based design. Circuit output is the Rluc luminescence over the Fluc luminescence (n=4); Crossbar, mean. **d**, Constructs used to test for cross-reactivity between Flp and B3 recombinases. **e**, Test of cross-reactivity between Flp and B3 recombinases. Circuit output is the Rluc luminescence over the Fluc luminescence (n=4); Crossbar, mean. **f**, Summary of cognate and non-cognate recombination activity of Flp and B3. Solid line represents confirmed recombination event while the dashed line represents a tested but not active recombination event. Some figure components created with BioRender.

To create a 2-input OR gate using the identify function, we repressed output gene expression with the *OCS* terminator, with subsequent removal by Flp and/or B3. Our OR gate has a single *OCS* terminator surrounded by recombination sites for both Flp and B3, so either recombinase can remove the terminator and activate output expression (Fig. 3a). Twenty four hours (Fig. 3b) and 48 hours (Fig. 3c) after protoplast transfection, expression of either recombinase activated the circuit’s output to levels at or above the constitutive expression reporter. Curiously, after 48 hours, some of the circuit state samples had even higher levels of reporter expression than the constitutive expression control, suggesting that the recombination site remaining after recombination in the 5’ UTR region can enhance the expression (Fig. 3c). To construct an AND gate, two *OCS* terminators were placed within the 5’ UTR, each with a different pair of recombination sites, so that each recombinase removes one, but not both terminators (Fig. 3e). The 1-input states remained repressed 24 hours after transfection, as expected, while the 2-input states were activated, but not to the same extent as the constitutive expression control (Fig. 3e). After 48 hours, the 2-input states were expressed at similar levels to the constitutive expression control (Fig. 3f), suggesting that activation of the AND gate, requiring two recombination events, was slower than the OR gate (Fig. 3e). We then constructed a single AND gate circuit that could be switched between its different states based on the activity of cell type-specific and chemically-inducible promoters driving the recombinase expression to test the circuit functionality *in vivo*. The cortex-specific *CO2* promoter^45^ was used to drive *B3*, while the dexamethasone (DEX)-inducible *pOp6* promoter^46^ drove *Flp* expression (Fig. 3g). Therefore, this AND gate would only activate the expression of nuclear localized GFP in cortex cells after DEX induction (Fig. 3g). We visualized transgenic Arabidopsis roots and observed strong GFP signal in the nuclei of cortex cells, but not in other root cell types, and only in the presence of DEX compared to DMSO solvent control (Fig. 3g). Together, this demonstrates construction of functional recombinase-based AND and OR gates that exhibit memory of the input signal in plant cells.

**Fig. 3:**
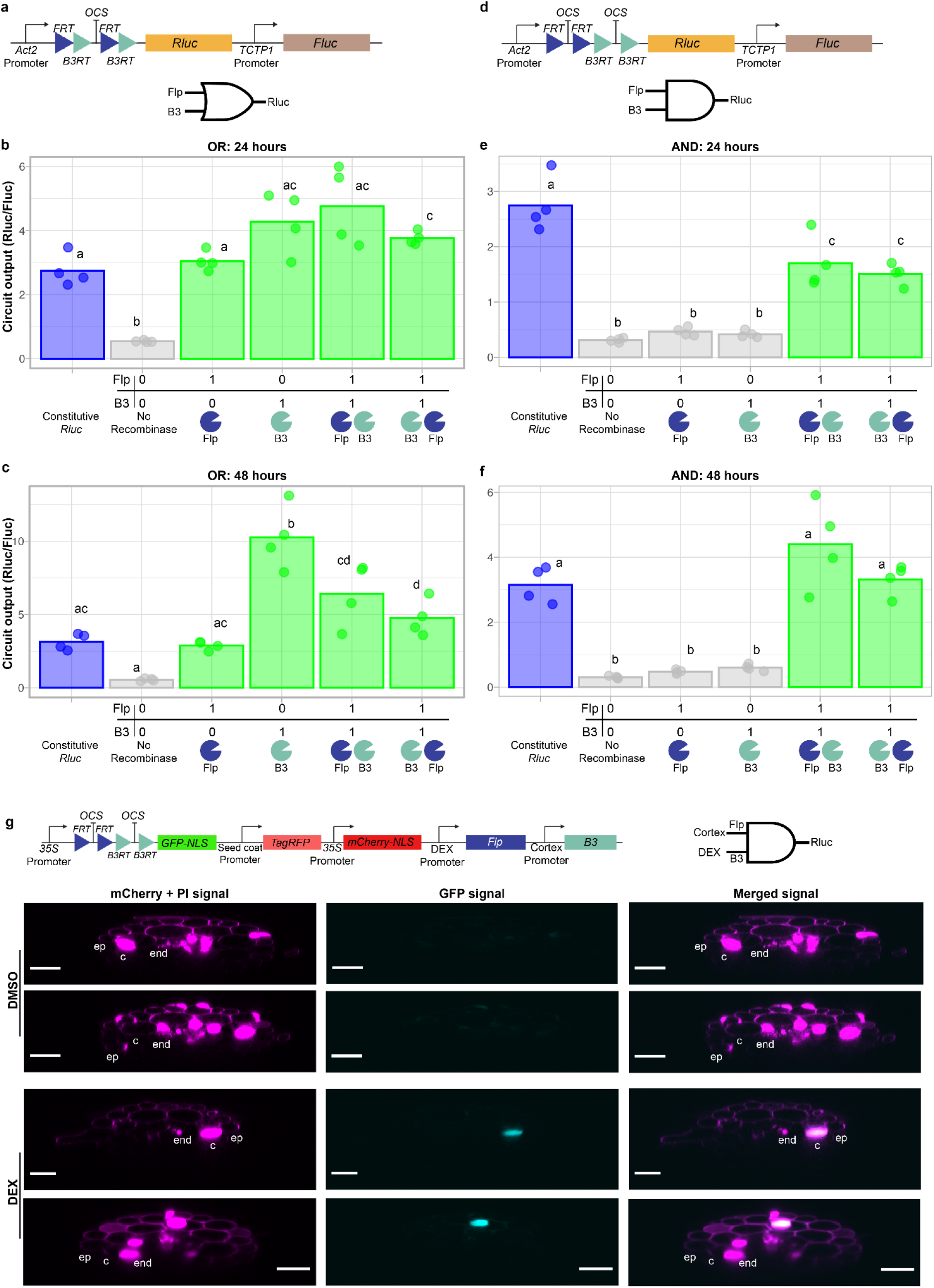
AND and OR gate construction using recombination-based gene circuits in plants. **a**, The output and normaliser units of the OR gate construct design. **b**,**c**, A 2-input OR gate that produces high circuit output activity when one of the two inputs is present, **(b)** 24 or **(c)** 48 hours after Arabidopsis protoplast transfection. Circuit output activity is the Rluc / Fluc luminescence ratio; also for panel d) (n=4); Crossbar, mean. Bar colours and letters representing statistical differences per Fig. 2. **d**, The output and normaliser units of the AND gate construct design. **e**,**f**, A 2-input AND gate that produces high Rluc luminescence relative to Fluc luminescence when both of the inputs are present, **(e)** 24 or **(f)** 48 hours after Arabidopsis protoplast transfection. Circuit output is the Rluc luminescence over the Fluc luminescence (n=4); Crossbar, mean. Bar colours as per Fig. 2. **g**, AND gate functionality *in vivo*, driven by condition-specific promoters. Schematic of the construct at top. Confocal images from roots of induced (dexamethasone[DEX]-treated) and uninduced (DMSO solvent control) T3 transgenic plants, with optical transverse sections of roots shown. Expression of the output gene (nuclear localizedlocalised GFP) is blocked unless in a cell with both recombinases expressed, where Flp expression is induced when exposed to DEX and B3 expression is driven by the cortex-specific *CO2* promoter. These plants also express nuclear localized mCherry from the *35S* promoter and were stained with the cell wall stain propidium iodide (PI). The magenta (mCherry and PI) represents the infocus nuclei and cell walls, while the cyan (GFP) represents the induced nuclei in the cortex. Two representative plants per condition. The outermost cell layer is the epidermis (ep), then the cortex (c), and the layer below that is the endodermis (end). Scale bar is 20 μm. Some figure components created with BioRender.

So far, all of our tested circuits have been based around activation (identify function), but the negation (NOT) function to repress a target gene is equally important in creating a fully programmable system. Therefore we designed a 2-input NOR gate that will repress the target gene when either of the inputs are present by flanking the output gene CDS by recombination sites (Supp. Fig. 3a). However, while Flp was able to repress Rluc output 24 hours after transfection, B3 was not (Supp. Fig. 3b). Furthermore, after 48 hours, though B3 did repress circuit output by 2-fold, this was to a lower extent than that achieved by Flp (10-fold reduction; Supp. Fig. 3c). It is possible that B3 is less efficient when the recombination sites are located further away from each other (∼2 kb) when flanking the CDS, than when they were used in the identify function circuits (∼0.7 kb apart). Therefore, we designed single input NOT gates to test each recombinase in isolation as well as a second generation of negation function circuits that remove the smaller *Act2* promoter (∼0.8 kb) instead of removing the large CDS, to test whether this will lead to faster activity state switching for B3. Twenty-four hours after transfection, the CDS-targeting Flp NOT gate repressed Rluc activity to ∼20% of the 0-input state (Fig. 4a; Supp. Fig. 4a), while the CDS-targeting B3 NOT gate did not repress the Rluc target (Fig 4a; Supp Fig 4a). However, both the Flp and B3 promoter-targeting NOT gates repressed Rluc activity to <25% after 24 hours (Fig. 4a; Supp. Fig. 4), suggesting that, at least for B3, the promoter targeting design leads to faster and more efficient negation function than targeting the CDS. Even the CDS-targeting B3 NOT design was able to repress circuit output to ∼25% relative to the 0-input control, but it took 64 hours to achieve a similar level of repression as the other designs could achieve within 24 hours (18-35%; Fig. 4a and b), supporting the hypothesis that B3 is able to remove the larger CDS region, but that it is slower than when targeting the shorter terminator sequence (Fig. 2) or the promoter (Fig. 4a and b). It was surprising to see variation in the behavior of these two recombinases in NOT circuits given that they performed similarly in the identify circuit designs (Fig. 2), which could allow for future customization of circuit timing. For all NOT gate designs tested, repression increased over time, suggesting that the 24 hour time-point underestimates the true extent of the difference between the on and off states. Knowing the optimal design of NOT logic for B3, we next designed and tested an A NIMPLY B logic gate (Fig. 4c), which combines identify and negation logic into one gate. Here, Flp is used to release the output from repression due to the *OCS* terminator, while B3 acts as a dominant repressor to block activation if B3 is expressed (Fig. 4c). To ensure sufficient time for the circuit to resolve, we measured the reporter genes at 64 hours post transfection and found that Flp was able to activate circuit output, while B3 blocked activation (Fig. 4c), thus demonstrating the expected A NIMPLY B logic. Taken together, we have demonstrated that the negation (NOT) function can be achieved in plant cells using recombinase targeting to create complex logic gates.

**Fig. 4:**
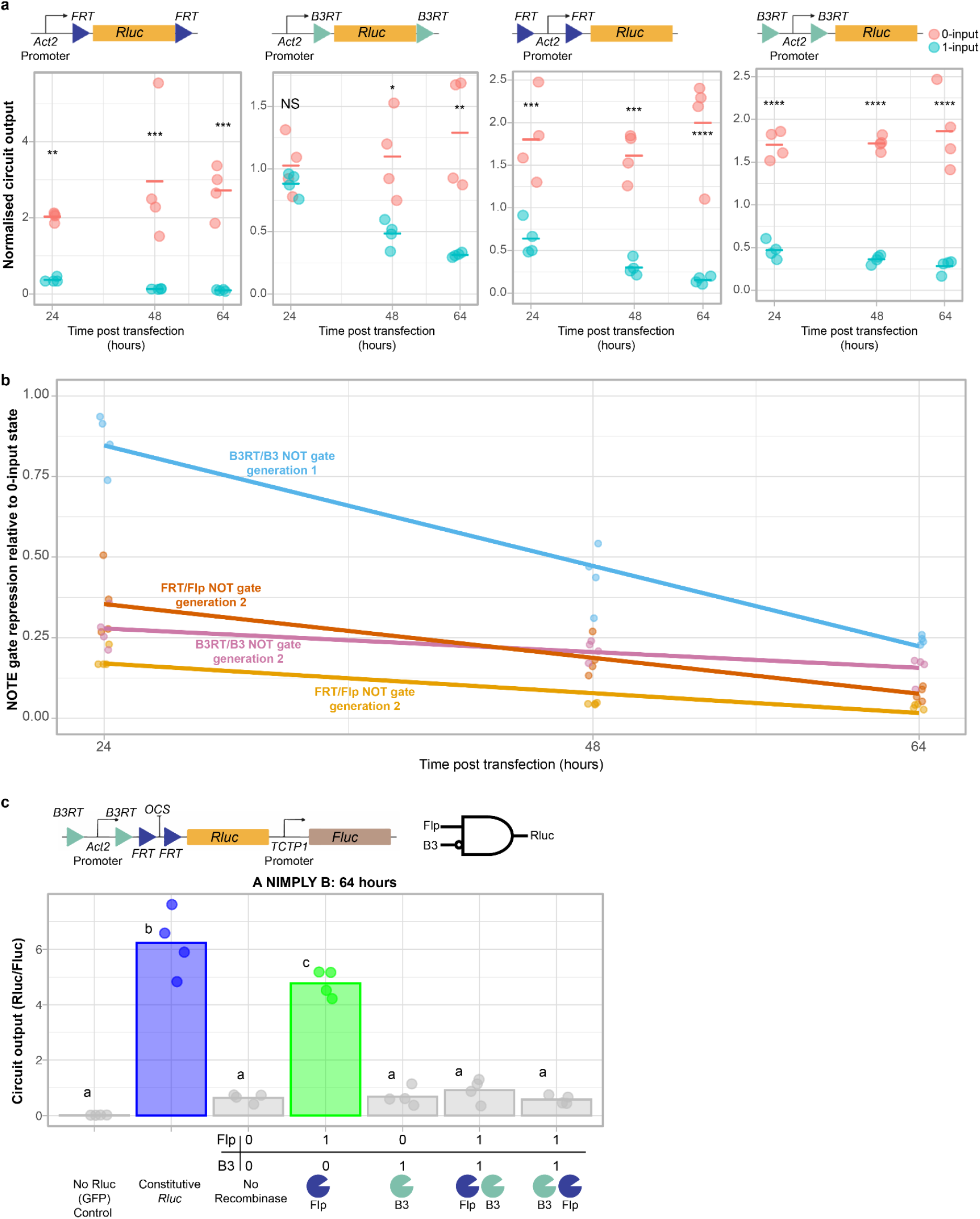
Negation (NOT) function implementation in plant cells by output gene CDS or promoter excision. **a**, Comparison of output (Rluc) repression over time with four different negation (NOT) function designs in plant cells. Right to left, Flp targeting the output gene CDS, B3 targeting the output gene CDS, Flp targeting the output gene promoter, and B3 targeting the output gene promoter. Three separate protoplast transfections were measured, one after 24 hours, one after 48 hours and one after 64 hours. Normalised circuit output is the Rluc / Fluc luminescence (circuit output) of the NOT function plasmid, divided by the circuit output of the Constitutive Rluc construct (Fig. 1b) to account differences between plates and time-points (n=3-4); Crossbar, mean. Asterisks denote significance from the ANOVA’s post-hoc test Tukey’s HSD: NS = P > 0.05, * = P ≤ 0.05, ** = P ≤ 0.01, *** = P ≤ 0.001, **** = P ≤ 0.0001. **b**, Circuit output of NOT function plasmids of the 1-input states (recombinase gene present) relative to the 0-input states (no recombinase gene present) over time. Generation 1 refers to NOT function designs where the recombinase targets the output gene CDS while generation 2 refers to the NOT function designs where the recombinase targets the output gene promoter. **c**, The A NIMPLY B gate combines the identify (activation) function and the negation (NOT) function and was achieved in plant cells by targeting the Flp recombinase to remove the repressive OCS terminator upstream of the output (Rluc) CDS, thus activating the circuit when only Flp was expressed, and the B3 recombinase to remove the Act2 promoter upstream of the output (Rluc), thus repressing the circuit regardless of the activity of Flp. Circuit output is the Rluc luminescence over the Fluc luminescence (n=4); Crossbar, mean. Bar colours as per Fig. 2. Bars with different letters have a significant difference calculated by a one-way ANOVA and Tukey’s HSD test (p < 0.05). Some figure components created with BioRender.

Thus far, the 2-input logic gates that we have demonstrated are asynchronous logic devices: they can be triggered by two stimuli in the same cell, but not necessarily occurring at the same time. Sometimes it may be beneficial to have the output triggered only when both inputs are occurring simultaneously, thereby adding an extra condition to the logic function. Thus we took advantage of the split-Flp recombinase, where both halves of the split-Flp protein are inactive and only allow for recombination after dimerisation through a separate dimerisation domain occurs^19,47^. To achieve this, we took the C1 homo-dimerisation domain from phage and fused it via a linker to each Flp fragment, given it was highly effective when used in a split-TALE system^24^. When only one Flp fragment-encoding gene was included on the plasmid, no activation of the repressed *Rluc* gene was observed (Fig. 5a). However, when both Flp fragments were expressed, Rluc activity was comparable to the intact *Flp* gene-encoding plasmid (Fig. 5a). This demonstrates that the C1 domain allows for highly efficient dimerisation and reconstitution of the Flp recombinase from fragments, enabling construction of a “split-AND” gate that is an example of a synchronous logic gate. Having demonstrated that the identify function works with the split-recombinase approach in plant cells, we then redesigned the construct to test the negate (NOT) function to create a “split-NAND” gate. As observed in Fig. 5, Rluc activity becomes repressed when both Flp fragments are present, indicating successful construction of a NAND gate.

**Fig. 5:**
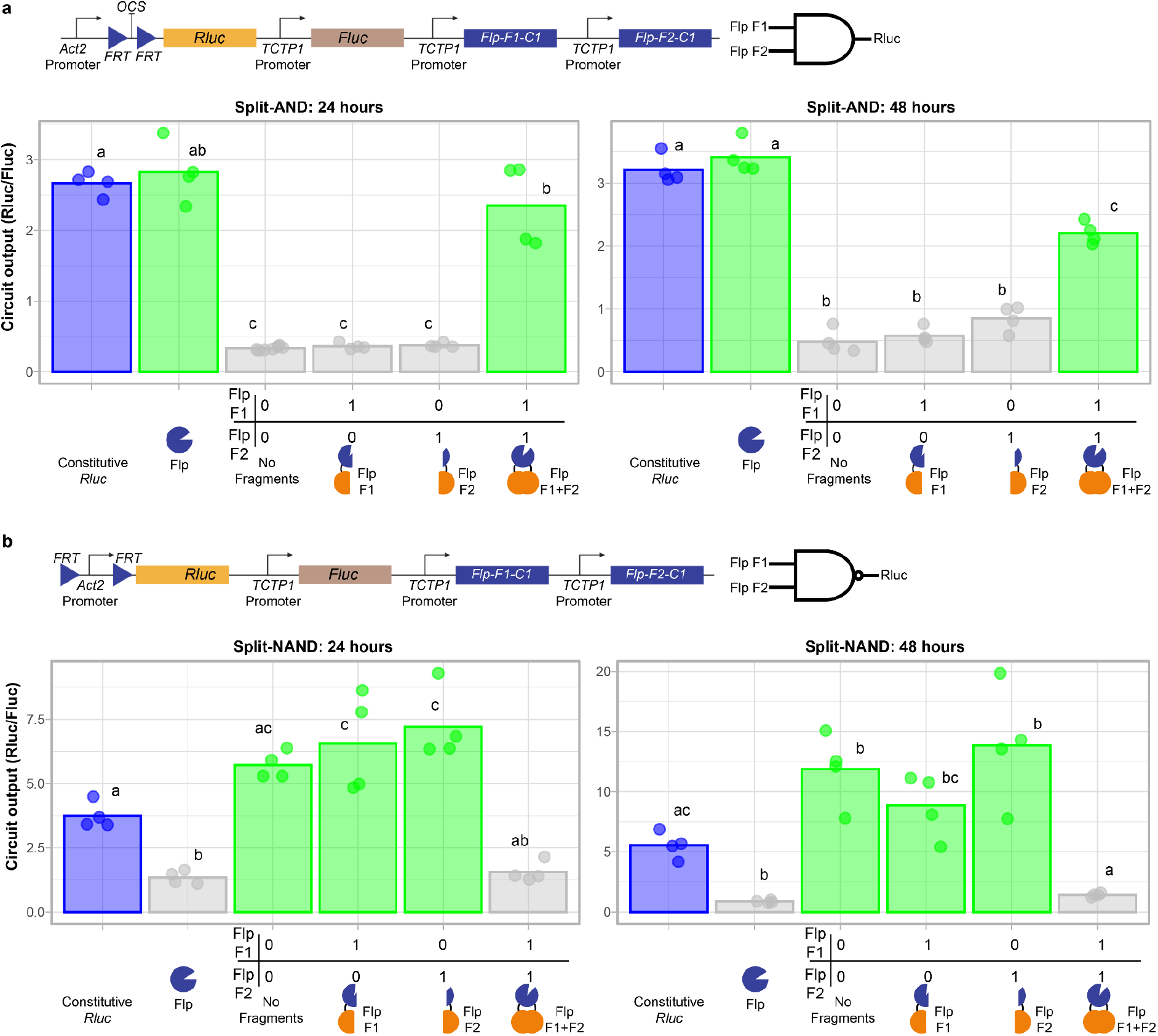
Construction of AND and NAND gates using split-recombinases. **a**, Flp split into two recombinase fragments (F1 and F2), each fused to the phage C1 dimerisation domain, to construct an AND gate through removal of an FRT-flanked OCS terminator located in the 5’UTR of the *Rluc* output gene, measured either 24 hours or 48 hours after protoplast transfection. Schematic represents the 2-input construct design. Crossbar, mean (n=4). Bar colours and letters representing statistical differences per Fig. 2. **b**, Flp split into two recombinase fragments (F1 and F2), each fused to the phage C1 dimerisation domain, to construct a NAND gate through removal of the FRT-flanked *Act2* promoter upstream of the *Rluc* output gene, measured either 24 hours or 48 hours after protoplast transfection. Schematic represents the 2-input construct design. Crossbar, mean (n=4). Bar colours and letters representing statistical differences per Fig. 2. Some figure components created with BioRender.

## Discussion

Here we have developed and implemented a broad suite of recombinase-based gene circuits and logic gates that demonstrate robust functionality in Arabidopsis protoplasts as well as transgenic plants. Using a high-throughput protoplast system, we identified effective components to construct Boolean logic gates in plant cells, and demonstrated that the Flp and B3 recombinases can be used to actualize a range of key 2-input Boolean logic, using both identify (activation) and negation (repression) functions. Furthermore, we have demonstrated that these circuits can be used to integrate two different condition-specific plant promoters to create a system that only induces expression in a specific cell type upon chemical treatment. The use of split-recombinases, where a single recombinase is rendered inactive unless both are expressed and can reconstitute into a functional dimer, can also be used to develop logic gates in plants, expanding the repertoire of components for constructing gene circuits. Here, we were able to directly engineer a whole plant with a gene circuit, and used endogenous signals (root cortex cell-type) and exogenous signals (chemical activation) to induce circuit output. Further advancement of gene circuits like these are needed to facilitate the development of more complex traits in transgenic crops, to help overcome the challenges of climate change, novel pathogen strains, and increased demand for food in both the field and controlled environments.

Expression of transgenes often needs tight control to ensure the fitness of the plant. For example, overexpression of oil producing enzymes in leaves can cause tissue destruction^3^, but delaying expression until senescence allows for oil to be accumulated^4^. Earlier induction when the leaf first matures would likely overcome the destruction associated with continuous expression and further increase yield. Continuous overexpression of a stress regulator can improve tolerance to drought^2^ but also negatively impacts overall plant growth^1,5^. By leveraging the sophistication of programmable gene circuits as reported here, an optimal balance could be achieved by tailoring activation of the circuit to only where and when it is needed to avoid the negative side-effects of overexpression.

Our study highlights some of the difficulties in developing new systems in synthetic biology. Even well-characterised genetic parts may not function as expected in new contexts. Of the six recombinases that we tested in our protoplast assay, only Flp and B3 were able to induce high levels of gene activation. This was unexpected given that Cre has been extensively used in plants before^32,44,48,49^. It may be possible that the protoplast system used to screen circuit components was not an optimal environment for Cre family recombinases. Alternatively, our constructs might not have been ideal for its expression in Arabidopsis.

Combining the induction properties of two promoters has previously been achieved in plants: root cell type-specific promoters were made DEX inducible by directly modifying the promoter sequences ^50^. Such a system is powerful but requires re-engineering of each promoter and is limited to only producing AND logic. Here, we present a system that can integrate the signals from two separate promoters and process them in a common processing unit, which can be further expanded with the integration of the demonstrated split recombinases. Our system also allows for full reprogrammability. All of the possible 2-input Boolean logic gates can be created with recombinase-based circuit components developed here, and we have demonstrated identify (activation: 1-input switches, OR, and AND gates) and negation (repression: NOT, NOR, and NAND) functions, and the A NIMPLY B gate that integrates both functions. Therefore, this recombinase-based gene circuit platform is highly flexible and programmable.

A recombinase-based gene circuit system comes with a number of unique features. Given that only a single recombination event per cell is needed to activate or repress the output, this offers a powerful and simple approach to transform a lowly expressed input signal, such as from a weak cell type-specific endogenous promoter, into a strong activation signal via the inherent activity of the recombined target promoter, taking advantage of the analog-to-digital conversion feature of recombinases^20^. This would also overcome the significant challenge of imbalanced input signal strength (expression) that can cause the logic of circuit designs based on transcriptional regulators to break down. One potential limitation with recombinase-based circuit design is the lack of reversibility. A different circuit design using artificial transcription factors could induce a reversible switch between states, as the inducing signal is removed. However, for many applications, a circuit that remembers the state it was switched to can be vital. For example, a light switch requiring continuous pressing for illumination would be highly inconvenient. The same would be true for a circuit in a crop that requires activation by an ongoing treatment signal in order to prepare it for an incoming stress. This is not unlike normal growth, where epigenetic marks can create semi-stable changes in response to internal developmental cues, or external environmental changes^26^. While epigenome editing tools are being developed, they often do not induce strong repression of the target, or create a stable memory, as occurs in natural epigenome regulatory processes^26^. Thus, until these limitations are overcome, plant gene circuits that feature stable memory via recombination will be particularly useful in a wide range of applications. In this study, we used tyrosine recombinases, but serine recombinases can create stable rearrangements that are reversible when a second factor is added. This has been used in plants to create a toggle switch, switching between two states (GFP vs RFP expressing)^25^. Further development of serine recombinases could overcome some of these limitations, and combination with the system presented here could allow for versatile and sophisticated gene circuits in the future. With characterization of additional recombinases that function effectively in plants, including more split-recombinases, this system could easily be expanded to create highly complex single layer gene circuits, integrating multiple inputs and potentially allowing for a single plant to respond to multiple environmental cues. Our diverse set of logic gates operating in plants with the ability to record a form of memory of past events has the potential to greatly advance plant biotechnology and complement transcriptional-based systems for gene circuits^7^.

## Materials and methods

### Plant materials and growth conditions

Protoplast experiments were performed using the Arabidopsis Col-0 strain. Arabidopsis plants for protoplast transfection and transformation via floral dip^51,52^ were grown on soil using a 16 hour light / 8 hour dark regime (22 °C / 18 °C). Plants for confocal analysis were grown on half Murashige and Skoog medium agar plates, with plates placed vertically in a growth chamber set to a 16 hour light / 8 hour dark regime (22 °C / 18 °C). All seeds were cold-treated at 4 °C for 2-4 days prior to transfer to the growth chamber.

### Plant transformation

We used the simplified preparation of Agrobacterium^52^ for floral dip transformation^51^ to generate stable Arabidopsis lines. Each construct contained a pFAST-R selection cassette^53^ from the MoClo plant parts kit^29^. Floral dipping into Arabidopsis transgenic line LhGR^46^ was performed to allow for activation of the DEX inducible gene circuit, and into Col-0 for the heat induced gene circuit.

Protoplasts were isolated from leaves with a modified Tape-Arabidopsis Sandwich protocol^33^, in which 10,000 protoplasts in 50 µl were added to each well for transfection in a 96-well plate with 5 µg of plasmid DNA (in 5 µl) before being mixed with 40% PEG and incubated for 15 minutes. Four replicates (wells) per construct per timepoint were transfected. Protoplasts were left in continuous light at 25 °C until the end of their incubation (24-72 hours later). The dual-luciferase assay (Promega) was then used to lyse the cells and measure Rluc and Fluc activity. Samples had a randomised placement in wells on the 96 well plate, to reduce confounding issues of positional effects on the plate.

### Plasmid construction

All plasmids were assembled into the backbones from the MoClo Plant Tool Kit ^29^, following the level 0, 1, and 2 system of parts to transcriptional units to multigene cassettes. These level 2 multigene cassettes were then used to transform plants, either via Agrobacterium-mediated floral dip to produce stable lines for *in vivo* work, or via PEG-mediated transient protoplast transfection. Plasmids were assembled in a one-pot restriction-ligation reaction^54^, in which 20-50 ng of uncut acceptor plasmid and a 2:1 molar ratio of donor plasmid or PCR products were mixed together with 1.5 μl DNA T4 ligase buffer (#B0202S; NEB), 0.0015 mg BSA (B9000S; NEB), 600 units of DNA T4 ligase (#M0202; NEB) and 5 units of *Bsa*I (#ER0291) or *Bpi*I (#ER1011) restriction enzyme (Thermofisher) in a total volume of 20 μl. The user-defined 4 bp overhangs produced by this reaction followed the MoClo and Phytobricks standards^29,55^. For cloning, plasmids were purified with the BIOLINE ISOLATE II Plasmid Mini Kit (#BIO-52057; Bioline). Before addition to the one-pot restriction-ligation reaction, DNA was purified by addition of 1 volume of 6 M guanidine hydrochloride followed by bead clean-up and elution in water. The 5’ UTRs containing recombination sites and repressive sequences (terminators or a uORF) were synthesized as two or three separate fragments as IDT gBlocks and assembled into one fragment into a level 0 ligation reaction with *Bpi*I cutting. The addition of the IV2 intron to the recombinase genes was performed by Gibson assembly. GeneWiz synthesized generation 1 FRT NOT/FER-B3RT NOR gates level 1 constructs.

### Plasmid-sequencing

Plasmids were sequenced after each stage of cloning via whole plasmid-sequencing (plasmid-seq), in which 100 ng of plasmid DNA was treated with Tn5 pre-loaded with sequencing adaptors, and incubated at 55 °C for 5-10 minutes. The tagmented DNA was added to a PCR reaction with 2x MyTaq mix (BIO-25041; Bioline) with primers that add library specific indexes and amplified for 10 cycles. The PCRs were then pooled together and bead cleaned, before being loaded onto a Illumina MiSeq to be sequenced (150 bp paired-end reads for at least 100x coverage). *De novo* assembly of the reads into a plasmid sequence was performed by Unicycler^56^ or aligned to a reference plasmid sequence with Bowtie 2^57^ to identify unexpected sequence changes.

### *in vitro* recombinase reaction

Isolated no recombinase OR gate plasmid (250 ng) was incubated with purified Cre recombinase (M0298S; NEB) at 37 °C for 1.5 hours followed by a 10-minute heat inactivation at 70 °C, after which a PCR was performed to amplify the region around the recombination target site.

### Confocal microscopy

Plants for DEX induction of cortex-specific GFP expression were grown on plates for ∼1 week before moving seedlings to a new plate with either 200 µM DEX (dissolved in DMSO) or equivalent concentration of DMSO and treated for 2-3 days before viewing on a Nikon C2+ confocal microscope. For heat-shock treated plants, ∼1 week old seedlings were moved to a 37 °C incubator (in the dark, wrapped in foil) for 5 hours, or left at room temperature as a control (in the dark, wrapped in foil). After heat-shock (or room temperature treatment), plants were returned to the growth room for 1-3 days before being taken to the microscope. GFP signal was collected in the FITC channel and the mCherry signal was collected in the TRITC channel, and transmitted light was also captured. For the heat-shock treated plants, a 10X objective was used to image the plants and in ImageJ, the Medium filter (2.0 pixels) was used to reduce noise. For the heat-shock experiment, the images are a merger of the transmitted light and the FITC (GFP) channels and used the Z-Stack Max Intensity projection in ImageJ from 10 slices. As for the DEX-induced plants, the 20X objective was used and in ImageJ the Medium filter (3.0 pixels) was used to reduce noise and the maximum brightness was set to 1000. The Z-stack was resliced from the “left” to view the roots as if looking up from the root tip and the representative images are a single slice from this view. Additionally for the DEX-induced roots, cell walls were stained with 10 µg/ml PI for 10 minutes in a dark room before being washed in water and imaged^58^.

### Figure preparation

Plots were generated in the R programming language with ggplot2^59^. Construct diagrams were created with Biorender.com. Figures were prepared in Adobe Illustrator.

## Acknowledgements

We would like to thank Brady Johnston for advice on R code, Marina Oliva and David Collings for help with confocal image analysis, Indra Roux for suggestion of the *in vitro* recombinase assay, Yit-Heng Chooi for the kind donation of purified recombinant Cre enzyme, Brian Crawford for the kind donation of a pOp6 promoter containing plasmid, Chris Helliwell for graciously providing the LhGR seeds used in this study, and Wilson Wong for generous donation of mammalian BLADE plasmids.

## Author Contribution

J.P.B.L. and R.L. conceived of the project and wrote the manuscript. J.P.B.L. with F.L. and P.G. conducted the experiments. J.P., T.S. and C.P. conducted the plasmid sequencing. M.A.K. developed the protoplast transformation protocol, and M.A.K. and B.K. provided technical assistance with the protoplast assay, luciferase assay, and cloning. All authors approved of and contributed to the final version of the manuscript.

