## Supplemental Figures for "Synthetic memory circuits for programmable cell reconfiguration in plants"

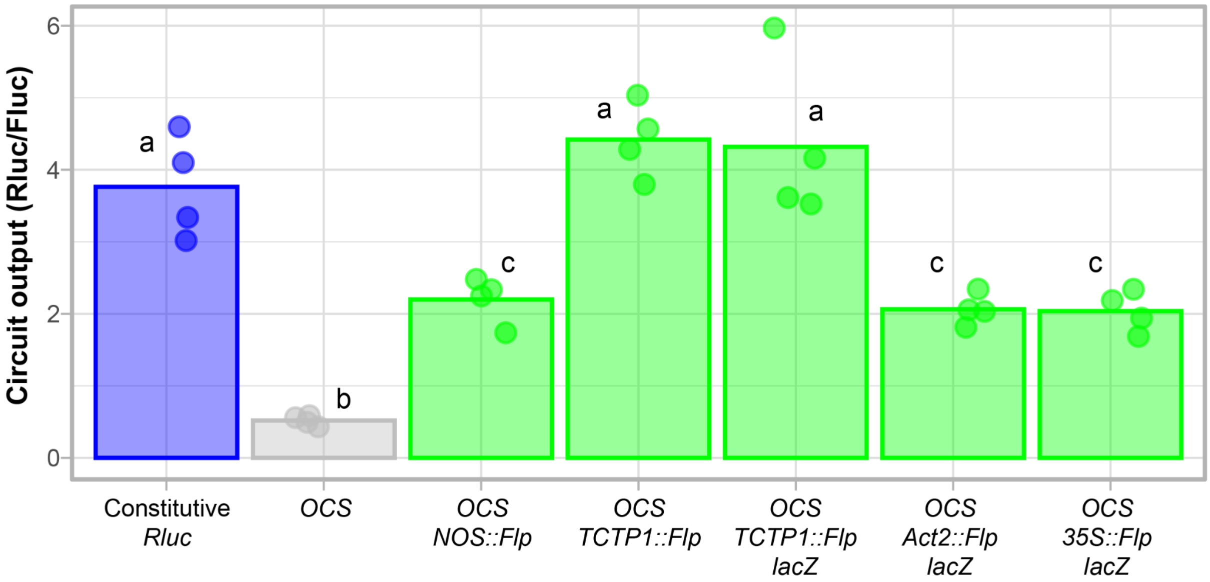


**Supplementary Fig. 1: Comparison of promoters drive recombinase for activation and effect of the presence of *lacZ* in the final construct**

Test of different promoters driving *Flp* recombinase expression in protoplasts (n=4). Crossbar, mean. Versions of the *TCTP1* promoter construct vary by presence/absence of the *lacZ* end linker [29,41] in the final construct, to aid in the selection of a positive clone. No difference between the variant with and without the *lacZ* included in the endlinker was observed. Blue bar is the control sample, green bars represent samples that are expected to be activated (turned on), gray bar represents the sample expected to be repressed (turned off). For this protoplast transfection experiment, circuit output activity is the Rluc / luminescence over the Fluc luminescence ratio, 24 hours after transfection (n=4). Bars with different letters have a significant difference calculated by a one-way ANOVA and Tukey's HSD test ($p < 0.05$).

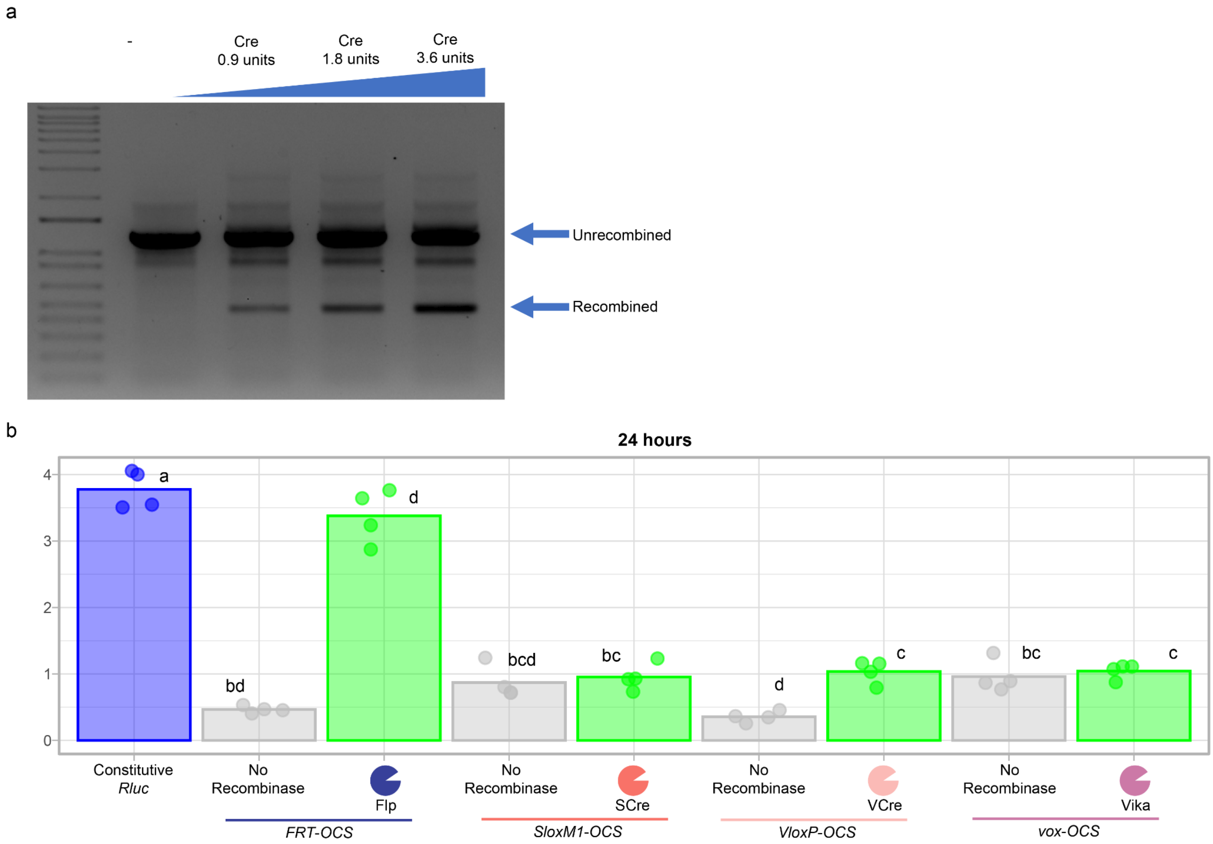


**Supplementary Fig. 2: Attempted Cre/Flp-based OR gate that resembled an A NIMPLY B gate**

**a,** *in vitro* recombinase assay of the OR gate. Isolated no recombinase OR gate plasmid was incubated with purified Cre recombinase and then amplified by PCR. With increasing amounts of Cre recombinase added to the reaction, the more recombined product is produced, showing that the plasmid can be recombined, even if we do not see activity in protoplasts.

**b,** Testing of alternative recombinases. SCre, VCre and Vika were tested for activation in an 1-input identify circuit design to identify a recombinase capable of activating the circuit to be used in a 2-input circuit. SCre (1.1x fold increase, $p = 1.0$) and Vika (1.1x fold increase, $p = 1.0$) did not activate target expression, while VCre had a modest level of activation (2.9 fold increase, $p = 0.006$), compared to Flp (7.3 fold increase, $p < 0.0000001$). Crossbar, mean. Blue bar represents the control sample, green bars represent samples expected to be activated (turned on), gray bars represent samples expected to be repressed (turned off). For this protoplast transfection experiment, circuit output activity is the Rluc / luminescence over the Fluc luminescence ratio, 24 hours after transfection (n=4). Bars with different letters have a significant difference calculated by a one-way ANOVA and Tukey’s HSD test ($p < 0.05$).

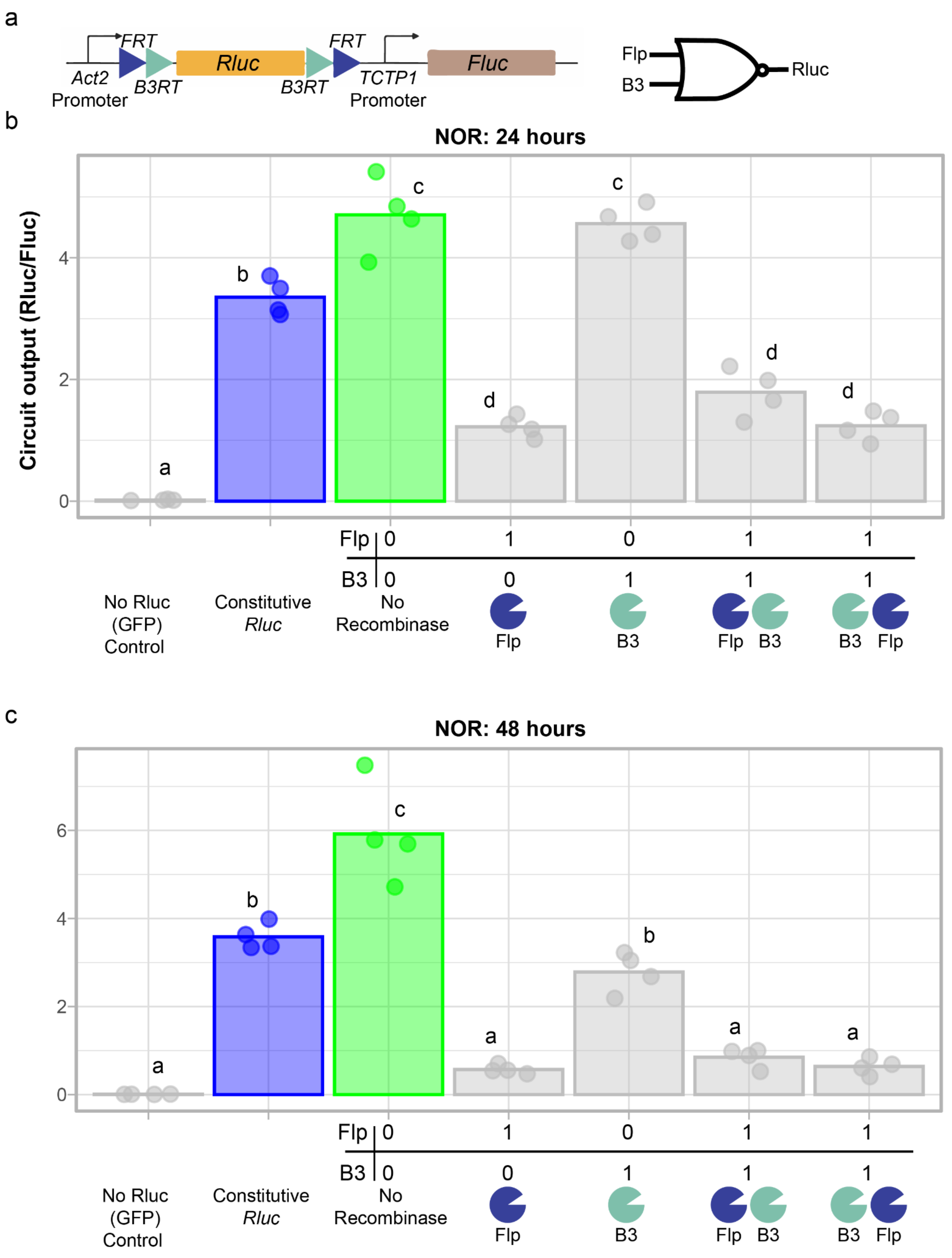


**Supplementary Fig. 3: A NOR gate with CDS targeting Flp and B3 recombinases has slow and modest switching when B3 is the sole input.**
**a,** The output and normaliser units of the NOR gate construct design.**b,** The NOR gate that produces high Rluc luminescence relative to Fluc luminescence only when Flp is at least one

of the inputs, 24 hours post-transfection. Circuit output is the Rluc luminescence over the Fluc luminescence (n=4); Crossbar, mean. Bar colours as per Fig. 2.
**c,** The NOR gate that produces high Rluc luminescence relative to Fluc luminescence when either input is active, 24 hours post-transfection. Circuit output is the Rluc luminescence over the Fluc luminescence (n=4); Crossbar, mean. Bar colours as per Fig. 2.
Bars with different letters have a significant difference calculated by a one-way ANOVA and Tukey's HSD test ($p < 0.05$).
Some figure components created with BioRender.

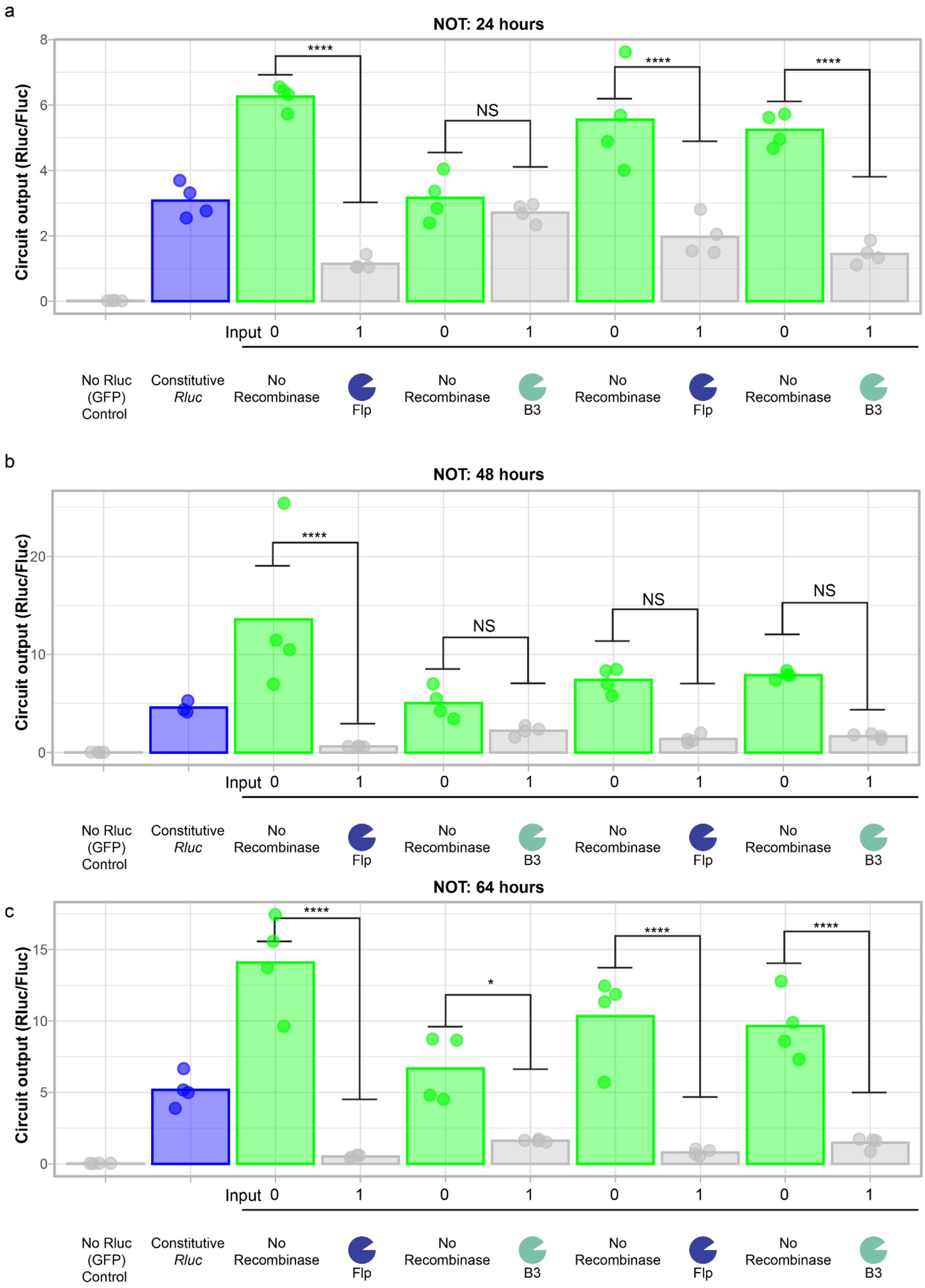


**Supplementary Fig. 4: Negation (NOT) function testing in protoplasts.**

**a,** Comparison of different negation (NOT) function construct designs, where Rluc luminescence relative to Fluc luminescence should be low when there is an input (recombinase) present, **(a)** 24, **(b)** 48, or **(c)** 64 hours post-transfection. Crossbar, mean. Gen (generation) 1 designs have the recombination sites flanking the output gene's CDS while gen 2 designs have the recombination sites flanking the output gene's promoter. Bar colours as per Fig. 2. For this protoplast transfection experiment, circuit output activity is the Rluc / luminescence over the Fluc luminescence ratio, 24 hours after transfection (n=3-4). Asterisks denote significance from the ANOVA's post-hoc test TukeyHSD: NS = $P > 0.05$, * = $P \leq 0.05$, ** = $P \leq 0.01$, *** = $P \leq 0.001$, **** = $P \leq 0.0001$.
